# Not So Mosaic After All? The Core Genomes of IncP-1 Plasmids Evolve Predominantly Through Vertical Transmission

**DOI:** 10.1101/2025.08.13.669769

**Authors:** David Sneddon, Clinton A. Elg, Jack Sullivan, Eva M. Top

## Abstract

Broad-host-range plasmids of the incompatibility group IncP-1 play an important role in bacterial evolution by disseminating genes between diverse bacteria. This includes the notable spread of antibiotic resistance genes to bacterial pathogens that increasingly threaten human health. A better understanding of the evolution of genetic elements like IncP-1 plasmids that underwrite bacterial adaptation is required to counter this threat. Here we examined the evolutionary history of IncP-1 plasmids by utilizing the large increase in published genomes and advancing techniques for lineage-based detection of microbial recombination. We specifically tested whether IncP-1 backbone genes evolve primarily *(1)* through vertical transmission, or *(2)* horizontally through homologous recombination between distinct plasmids. Our key finding is the IncP-1 plasmid backbone evolves predominantly through vertical transmission, which was supported by a nucleotide-level mapping of backbone recombination. We also showed via a phylogeny of 187 IncP-1 plasmids that established subgroup categories are well-supported.

Finally, we examined factors that are potentially associated with IncP-1 subgroup diversification, including a gene related to plasmid-host adaptation and accessory genes like antibiotic and resistance metals. While some IncP-1 subgroups do correlate with certain categories of accessory genes, it remains unknown if these are causes or consequences of subgroup diversification.

## Introduction

Self-transmissible broad-host range (BHR) plasmids are known for their ability to transfer genetic material horizontally between diverse species of bacteria via cell-to-cell transfer called conjugation (Datta and Hedges 1972; Frost et al. 2005; Norman et al. 2009). This horizontal transfer of genes is a key driver of bacterial evolution by allowing bacteria to rapidly acquire new traits, such as virulence factors, catabolic functions, or heavy metal and antibiotic resistance (Goldenfeld and Woese 2007; Soucy et al. 2015; Hall et al. 2017). For example, the rapid spread of antibiotic resistance genes carried on plasmids can lead to the emergence of multidrug-resistant (MDR) pathogens, posing significant challenges for human health (Partridge et al. 2018). Despite the threat of plasmids disseminating MDR and virulence factors (Pilla and Tang 2018), relatively little is known about the evolution of plasmids and its effect on the threats posed by pathogenic bacteria.

The IncP-1 group of plasmids stands out for a broad host range reflected in their ability to transfer between and replicate within diverse species of bacteria (Thomas and Smith 1987). IncP-1 plasmids tested to date can transfer into, establish and maintain themselves within multiple classes of Proteobacteria (Olsen and Shipley 1973; Schmidhauser and Helinski 1985; De Gelder et al. 2007; Shintani et al. 2010; Yano et al. 2012; 2013; Klümper et al. 2015; Kottara et al. 2016). They have even been shown to mobilize genes into Gram-positive bacteria (Musovic et al. 2006) and eukaryotic microbes (Heinemann and Sprague 1989). The transfer, replication, and maintenance functions needed to benefit these plasmids are encoded by a region of ‘backbone’ genes that share homology across IncP-1 plasmids. These backbone genes represent the core genomes of this plasmid group. In addition, IncP-1 plasmids often contain genes that encode traits beneficial to their host bacteria including MDR, catabolic pathways, and heavy metal resistance (Schlüter et al. 2007; Sen et al. 2011; Popowska and Krawczyk-Balska 2013). Such ‘accessory’ cargo genes are highly variable across IncP-1 plasmids and are almost always encoded in two specific accessory gene regions (Sota et al. 2007). Thus, the combined capabilities of IncP-1 plasmids to rapidly acquire and disseminate resistance and other beneficial genes across diverse taxa of bacteria contribute to their role as potent vehicles for bacterial adaptation (Popowska and Krawczyk-Balska 2013).

While the contributions of plasmids like the IncP-1 group to bacterial evolution is increasingly appreciated, our understanding of the evolutionary history of the plasmids themselves remains problematic. There are two possible models for evolution of the IncP-1 plasmid backbone, which we describe as horizontal or vertical. Under the horizontal model, plasmid backbone genes are thought to have different evolutionary histories due to frequent homologous recombination between similar plasmids. This contrasts with the vertical model, in which genes within a plasmid backbone are expected to share the same evolutionary history due to vertical inheritance. These different models have important implications for how IncP-1 plasmids evolve.

IncP-1 plasmids are historically classified into subgroups α, β, γ, δ, ε, ζ, η, θ, κ, ι, ο, μ, λ, ρ (Chikami et al. 1985; Pansegrau et al. 1994; Thorsted et al. 1998; Vedler et al. 2004; Norberg et al. 2011; Haines et al. 2006; Brown et al. 2013; Hayakawa et al. 2022a; Bahl et al. 2007). These classifications are based on overall sequence similarity of sister plasmids, as well as monophyly within phylogenies of individual or few backbone genes. However, these phylogenetic inferences prove troublesome as individual plasmid backbone genes that are putatively inherited via homologous recombination from different ancestral plasmids can significantly bias our estimation of evolutionary history when sampling of loci and plasmids are limited. (Schlüter 2003). The literature presents different findings about the degree of a horizontal versus vertical evolutionary history of the IncP-1 plasmids backbone. While one study concluded horizontal recombination is a prominent feature of IncP-1 backbone evolution (Norberg et al. 2011), another suggested a vertical evolutionary history based on successful reconstruction of resolved phylogeny (Sen et al. 2013). There remains a need to understand whether the IncP-1 plasmid backbone predominately evolves through mutation during vertical inheritance or horizontally through recombination. This will help elucidate the host range of these plasmids, and thereby the host trajectories through which their genetic traits like antibiotic resistance spread within microbial communities.

Sen et al. (2013) recovered four major topologies across 21 backbone genes, one of which was supported by 46% of gene trees. This topology was deemed to be the “true tree,” although a single concatenated phylogenetic analysis produced a different topology. This discordance was due to the weighting of an alternative topology represented by three genes, including one disproportionately large gene, *traE*. Here, we expand upon these previous studies by including many more individual plasmids, as well as employing robust methods that are not as susceptible to biases due to discordant evolutionary histories represented in large loci (Zhang et al. 2018).

We further explore the presence of and quantify homologous recombination between recognized subgroups of the IncP-1 group of plasmids using techniques for lineage-based detection of microbial recombination (Mostowy et al. 2017). Our key finding is that vertical inheritance is the predominant factor explaining the evolutionary history of IncP-1 plasmid backbones. This conclusion is based on the well-resolved phylogeny of 187 IncP-1 plasmids,as well as a detailed nucleotide-level mapping of recombination among 20 concatenated backbone genes. Our robust phylogeny showed that established IncP-1 subgroups correspond well to monophyletic groups (*i*.*e*., they descended from a common ancestor) with detectable vertical history. While our analysis of homologous recombination did identify examples of recombined backbone genes, including several that were previously discovered (Schlüter et al. 2007; Norberg et al. 2011), we found such recombination is comparatively rare (~1.4% of DNA) within the large set of IncP-1 plasmids we examined. Thus, both the phylogenetic and recombination analyses provide clear evidence for the predominance of vertical inheritance in the evolution of IncP-1 plasmid backbones.

Given that the existing IncP-1 subgroups are genetically distinct through vertical inheritance, we asked what may contribute to the diversification of these subgroups. For example, the bacterial host or the host’s environment might associate with subgroups. We took advantage of our curated IncP-1 dataset to analyze how genetic factors correlate with subgroups, including a known host-range factor and classes of genes including antibiotic resistance, metal resistance, and efflux pumps.

## Results

### Established IncP-1 subgroups are phylogenetically well-supported

We tested if an expanded set of IncP-1 plasmids would produce a well-supported phylogeny indicative of a vertical evolutionary history. Twenty backbone genes were identified to be present in all the plasmids within our dataset and were thereby selected for the analysis. This set included representative genes from each of the four regions of the IncP-1 backbone that collectively encode the core functions of IncP-1 plasmids (red, yellow, green, and blue in Figure 1).

**Figure 1.**
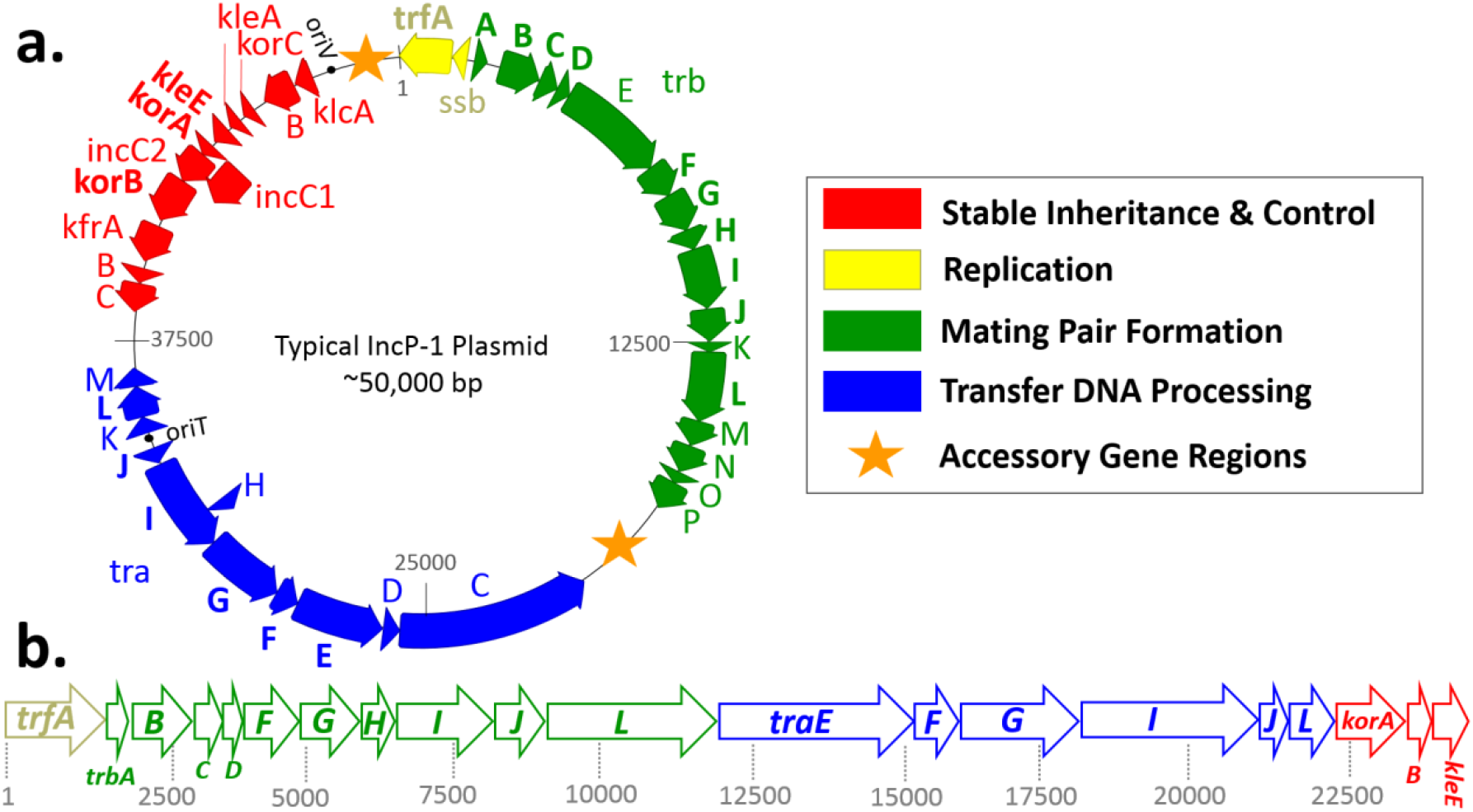
Genomic map of a typical IncP-1 plasmid and the backbone genes used in this study. **(a)** The four regions of the IncP-1 backbone are shown in red, yellow, green and blue (see legend). An orange star marks each of the two typical accessory gene regions. Bold text signifies backbone genes used in this study. **(b)** A map of the twenty concatenated backbone genes utilized in our lineage analysis.

Among our 20 gene trees we observed expected discordance, with all four gene tree topologies from Sen et al. (2013) represented within our set of loci, as well as a few additional topologies (Supplemental Data S1). The topology selected as the “true” tree in Sen et al. (2013) was represented in 11 of our 20 genes (Supplemental Data S2). This topology was also consistent with our summary lineage tree (Figure 2). Our lineage tree displayed monophyletic resolutions of all historically recognized subgroups with a few exceptions, supporting the hypothesis that all members within most subgroups descended from a common ancestor. All subgroup clades were well supported, with nodal support of ≥ 0.99 posterior probability. High nodal support was present across all deep nodes that define relationships among subgroups except for the nodes that unite IncP-1δ, IncP-1α, and IncP1-ε, in which moderate nodal support was observed (X=0.7, 0.59, 0.71, respectively). This well-supported lineage tree provides evidence of a strong signal of vertical evolutionary history among the 20 backbone genes we retained.

**Figure 2.**
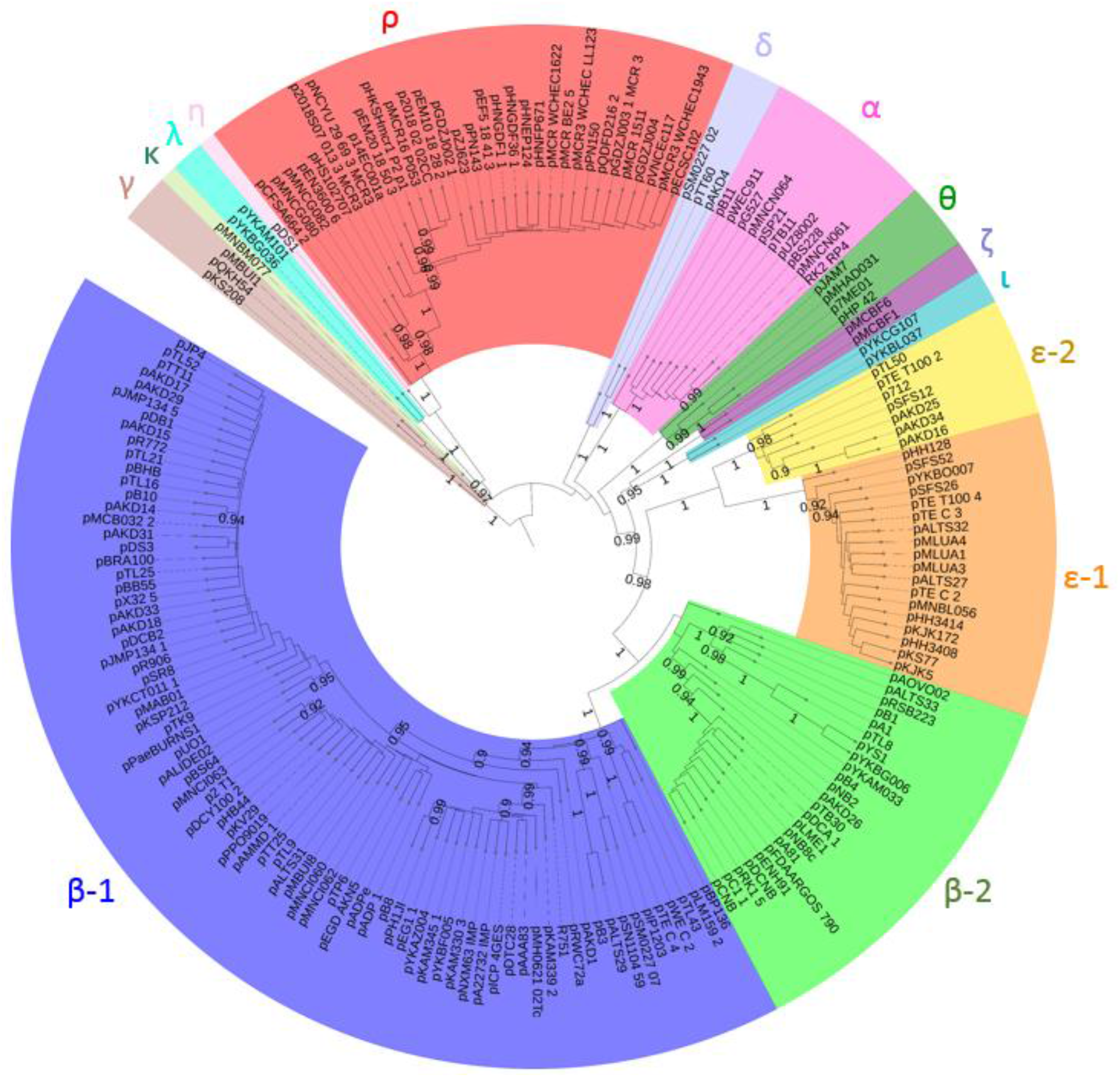
A summary lineage tree of 187 IncP-1 plasmids based on 20 backbone genes. The lineage tree shows monophyletic groups largely in agreement with established IncP-1 subgroups (organized by color). All posterior probabilities > 0.90 are annotated, which demonstrates strong nodal support for the initial branching/diversification, as well as monophyly of distinct subgroups.

The individual plasmids present in the monophyletic groups of our tree closely agreed with the 14 currently described IncP-1 subgroups: α (alpha; Chikami et al. 1985; Pansegrau et al. 1994), β (beta; Chikami et al. 1985; Thorsted et al. 1998), γ (gamma; Haines et al. 2006; Hill et al. 1992), δ (delta; Vedler et al. 2004), ε (epsilon; Bahl et al. 2007), ζ (zeta; Norberg et al. 2011), η (eta; Brown et al. 2013), θ (theta; Yakimov et al. 2016), and a 2022 study (Hayakawa et al. 2022b) added ι (iota), κ (kappa), λ (lambda), μ (mu), ο (omicron), and ρ (rho). The β and ε subgroups were further divided into β-1 and β-2 (Sen et al. 2010), and ε-1 and ε-2 (Bahl et al. 2009), respectively, and are each monophyletic within their respective subgroup. These subgroup divisions are displayed in Figure 2 as independent subgroups to display their monophyly.

Our expanded phylogeny suggests a reclassification of two recently proposed subgroups (Hayakawa et al. 2022b), which would reduce the number of subgroups from fourteen to twelve. The proposed ο subgroup contains a single plasmid, pYKBG036, which we find is sister with the plasmid from proposed subgroup λ, pYKAM101. We thus propose combining these two plasmids into λ, as the splitting into distinct subgroups does not improve monophyly. Next, the μ group plasmid pMNCG082 renders the ρ subgroup paraphyletic. Thus, we recommend adding pMNCG082 to the ρ subgroup. The net effect of these suggested changes is the merging of subgroup ο into λ, and subgroup μ into ρ, leaving twelve subgroups: α, β-1/2, γ, δ, ε-1/2, ζ, η, θ, ι, κ, λ, and ρ.

### Recombination of IncP-1 plasmid backbones genes is rare

We next sought to resolve the discrepancy between our lineage tree suggesting a vertical history and previous findings of horizontal recombination in regions of the IncP-1 backbone. In order to assess to what extent horizontal recombination is detectable in the evolutionary history of these IncP-1plasmid segments, we performed a search for recombined regions using a hidden markov model implemented in the software fastGEAR (Mostowy et al. 2017); see Methods). In this analysis, the nucleotide frequency within a subgroup represents a unique genetic signal. Every site of a considered plasmids backbone region was scanned and assigned to the subgroup whose genetic signal it best matches. Under a vertical evolutionary history, the plasmid backbone region will best match the genetic signal of its respective subgroup. Alternatively, under horizontal evolution the recombined regions of a plasmid backbone will match the signal of the subgroup with which recombination occurred. This approach allowed for an increased level of precision in detecting recombination within and between 20 backbone genes across 187 IncP-1 plasmids. As in the lineage tree analysis, the β-1/2 and ε-1/2 divisions were treated as independent subgroups.

We found that homologous recombination among IncP-1 plasmid backbone genes is rare. The most notable result was the degree to which vertical history is conserved in these 20 IncP-1 plasmid backbone genes. This was evident in the recombination visualization of Figure 3, in which small patches of between-subgroup recombination contrast with the overall pattern of vertical evolution of IncP-1 plasmid backbone genes. A total of 35 recombined regions were detected, 28 of which are unique to a single plasmid. The seven remaining recombined regions were conserved within lineages, each suggesting an ancestral recombination event that became vertically inherited. Most notable were 22 β-1 plasmids that have identical portions of *korB* and *kleE* from β-2 plasmids (Figure 3 bottom right-hand corner, in light green). To quantify the prevalence of recombination, we counted 65,369 base pairs as originating from external subgroups. The total number of base pairs considered was 4,575,516 (concatenated genes with base pair length of 24,468 × 187 plasmids analyzed). This suggests that only 1.4% of the analyzed IncP-1 plasmid backbone DNA had a detectable horizontal signal. Thus, 98.6% of the analyzed backbone genes had a vertical evolutionary history while less than 2% was attributable to horizontal between-plasmid recombination (Figure 3; for full resolution image, see Supplemental Data S3).

**Figure 3.**
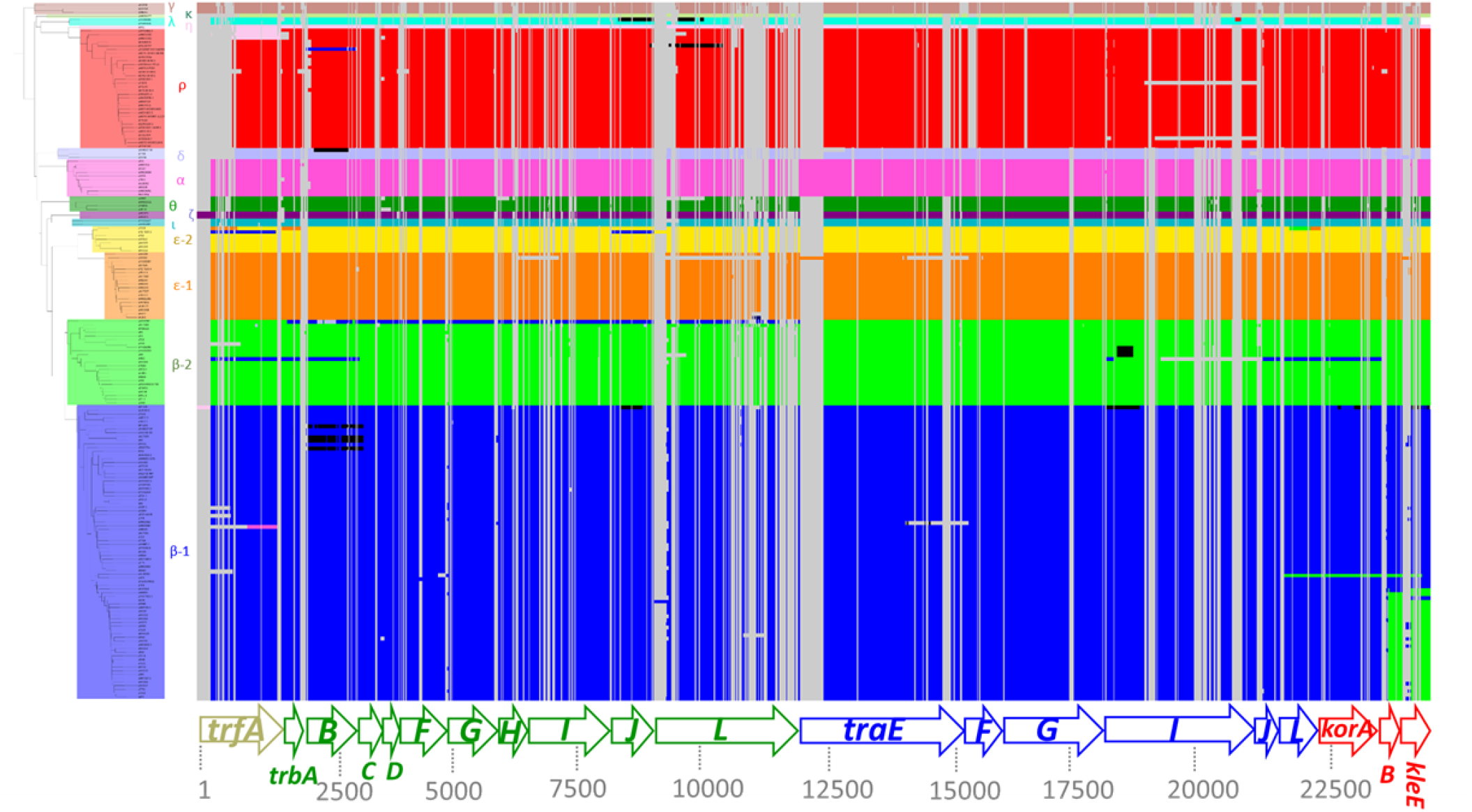
A map of backbone gene recombination between IncP-1 subgroups. Concatenated backbone genes (at bottom) give a nucleotide-level resolution of recombination detected in individual plasmid backbones (phylogenetic tips on left). Vertical grey bars represent gaps in alignment sequences. All other colors represent the genetic signal found within a respective subgroup. Under vertical transmission, the genetic signal of a plasmid’s genes will best match that of its subgroup (represented by matching colors). When a region of plasmid backbone is more similar to the genetic signal of a different subgroup, recombination is detected and colored with the plasmid subgroup with which it shares genetic similarity. Black represents regions with sections matching more than one subgroup.

To establish the accuracy of our recombination predictions we compared the exact genomic locations of detected recombination to previously reported IncP-1 recombination events. We found four specific examples of IncP-1 backbone region recombination in the literature: plasmids pB10 (Schluter 2003), pIJB1 (Sen et al. 2010), pB3, pBP136, and pAOVO02 (all from (Norberg et al. 2011). Plasmid pIJB1 is an unusual case of a repeated backbone region and was not included in our study. Plasmid pB10 was the first plasmid in which IncP-1 backbone gene recombination was detected, and our results likewise detected recombination in *kleE* (Schlüter 2003). The recombinant *trbB* previously identified in pB3 was present in our recombination data (Schlüter 2003). Norberg et al. (2011) previously predicted an undefined recombination in pAOVO02, and our analysis now provides precision by identifying *trbB* as the site of recombination. We also detected a recombined *trbJ* and *traJ* in pBP13632. In summary, the regions of orthology between subgroups as detected by fastGEAR support and inform the examples of IncP-1 plasmid backbone recombination identified in previous studies. The previously identified recombination was identified and represented in individual gene phylogenies by fastGEAR. The greatly expanded number of plasmids considered here and the increased resolution of our recombination analysis allowed us to demonstrate that recombination between IncP-1 plasmid backbones is rare (<2% of analyzed base pairs).

### Factors associated with IncP-1 subgroup diversification

The IncP-1 subgroups have distinct vertically inherited backbones, but what genetic factors might influence diversification into these subgroups? One possible factor could be plasmid adaptation to bacterial hosts, which has been observed in experimental evolution studies.

Additional factors that may contribute to IncP-1 diversification are the accessory genes. Plasmids containing accessory genes that are only advantageous to their host in specific environments may themselves become partitioned within that environment and thus become isolated from other IncP-1 plasmids that do not encode those host beneficial genes.

Searching the MEGAres (Bonin et al. 2023) database against our IncP-1 dataset yielded 401 antibiotic drug resistance, 79 metal resistance, and 54 multi-compound resistance genes (Supplemental Data S4). There was an average of 2.1 drug resistance, 0.42 metals resistance, and 0.29 multi-compound genes per plasmid. For each gene category, we took the average number of genes found within a subgroup and divided it by the average number of genes for all plasmids. The results are presented in Figure 4, where a value of 1 means that the subgroup has an average number of category genes, whereas a value of 2 shows this subgroup has on average twice as many category genes compared to all plasmids, and so on. A value of zero represents that class of genes was not found in a particular subgroup. Figure 4 shows clear differential patterns of accessory gene categories across subgroups. Of particular interest is the division of the ε subgroup: ε-1 plasmids have on average ~> 3-fold more drug and multi-compound genes than average, while ε-2 had ~0.5 the average number of hits in the same categories. This is consistent with the finding of Li et al. (2016) that in contrast to the IncP-1ε-1 plasmids, the ε-2 plasmids known at the time did not contain antibiotic resistance genes. Another interesting observation is the slightly higher gene content for all three categories in the β-1 group compared to β-2. This is consistent with a large clade of β-2 plasmids being catabolic plasmids that do not encode antibiotic resistance genes (Król et al. 2012).

**Figure 4.**
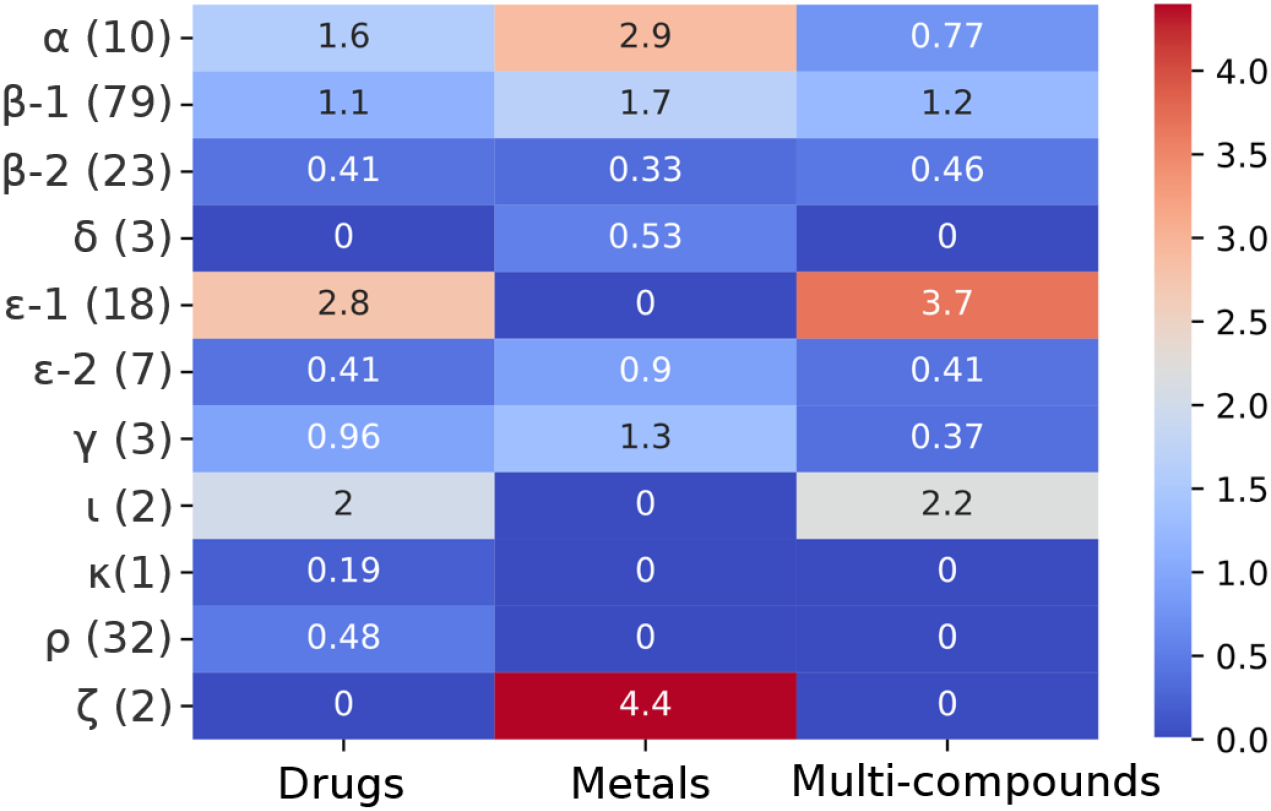
The relative frequency of classes of accessory region genes in 187 IncP-1 plasmids. Accessory regions genes were divided into sets of genes conferring resistance to drugs, or metals, or ‘multi-compound’ genes like efflux pumps which may confer some resistance to both (horizontal axis). Subgroups are on the vertical axis, with the parenthesis containing the sum of plasmids in that subgroup that contain at least one of the classes of genes examined. The numbers on the right-side scale are calculated from the average number of genes within a subgroup divided by the average number of genes across all plasmids.

## Discussion

Plasmids represent a molecular cornerstone, allowing rapid bacterial adaptation to changing environments, and are of particular importance in conferring MDR to pathogens that threaten human health (Castañeda-Barba et al. 2024). A better understanding of the evolution of IncP-1 plasmids, which can broadly disseminate such resistances between diverse species (Popowska and Krawczyk-Balska 2013b), is therefore of importance. We found that IncP-1 plasmid backbones primarily evolved through vertical transmission and were able to quantify the low prevalence and genetic location of horizontal recombination among a subset of backbone genes. Of particular interest are the factors that may lead to subgroup diversification (Li et al. 2016), as understanding the dynamics of IncP-1 plasmid diversification can provide crucial context to their role in bacterial adaptation, including the spread of genetic traits like MDR within bacterial communities.

Our lineage tree of 187 IncP-1 plasmids leveraged the increase of available plasmid sequences to provide new insights into their evolutionary history. Overall, we observed a clear vertical history of evolution represented in our individual-level phylogeny summarized across 20 plasmid backbone genes. The topology of our summary tree displays the same topology represented by the majority of our genes (11 genes; Supplemental Data S2); however, we do recover discordant topologies across other genes within our dataset. This signal is represented in the deep nodal support of our tree, in which discordant nodes that unite certain subgroups have significantly lower support than other deep nodes. In all but a select few cases as illustrated by our recombination analysis, IncP-1 subgroups are monophyletic regardless of the deeper resolution between subgroups. This indicates that this discordance among gene trees is likely caused by rare single recombination events deep in the tree that are vertically inherited by all plasmids in the recipient subgroup, or random sorting of ancestral alleles (incomplete lineage sorting).

As observed by Sen et al. (2013), the majority of individual gene tree topologies (11 of 20) support a single underlying topology in which IncP-1δ, IncP-1α, and IncP-1ε are all paraphyletic with respect to each other, and IncP-1ε is sister to IncP-1β (Topology 1 in Sen et al. 2013). The second most represented gene tree topology places the α subgroup sister to δ (5 genes). In previous studies similar to ours, large loci (e.g., *traE*) that may be involved in backbone recombination have biased overall phylogenetic signal towards this alternative topology, which prompted the removal of these loci from their analyses. Our summary-based phylogenomic approach is more robust to these alternative topologies, as all genes are weighted equally regardless of sequence length. This allows us to use an expanded set of genes, thus retaining any signal of inter-subgroup recombination deep in the tree.

Interestingly, the recent addition of newer subgroups (ρ, θ, ζ, ι, λ, η, κ) does not disrupt the overall topology recovered by Sen et al (2013) as the ρ, η, κ, and λ all form a monophyletic group and are sister to the rest of the IncP-1 ingroup. Subgroups ι and ζ are sister but are not monophyletic with any other individual subgroups. This is the case for θ as well. Across individual gene trees, resolution of κ, which is represented by a single plasmid (pMNBM077) is highly discordant. This could be due to inter-subgroup recombination that is not represented in our fastGEAR output. As this plasmid is the only representative of this subgroup, and is highly dissimilar to any other subgroup, our analytical framework lacks the statistical power to detect any recombination. To explore the recombination history of this plasmid under the framework presented here, more individuals belonging to the κ subgroup must be sampled.

We are indebted to the early work of previous studies that explored the evolution of IncP-1 plasmid backbone genes, often finding cases of recombination by painstaking manual analysis (Schluter 2003; Norberg et al. 2011; Sen et al. 2013). It was striking to find four cases where previous predictions matched our HMM predicted recombined regions (in pB3, pB10, pAOVO02, pBP136). These studies were limited at the time by the number of IncP-1 genomes available (ranging between 25 and 65), and the less precise methods of detecting recombination. For example, utilizing the pairwise homoplasy index (PHI; Bruen et al. 2006) allowed for statistical testing of backbone regions for recombination (Norberg et al. 2011), but this could not provide any genetic context on where or to what degree regions of recombination exist.

Additionally, only four subgroups had enough representative plasmids to be examined for recombination, compared to the twelve subgroups we analyzed here. Remarkably, six of these subgroups have been defined in just the last couple of years (Hayakawa et al. 2022). Our present work considering 187 plasmids and the nucleotide by nucleotide fine-resolution of recombination regions also agrees with the previous finding of a detectable vertical history in IncP-1 backbone genes (Sen et al. 2013). The low frequency and limited cases of backbone recombination found in our expanded study suggest that IncP-1 plasmid backbones primarily evolve by mutations that are vertically inherited, rather than by homologous recombination between plasmids.

Thus, it becomes apparent that the distinct IncP-1 subgroups generally display “recombination isolation,” which raises important questions about the host range, ecology, and evolution of IncP-1 plasmids. Are IncP-1 plasmid subgroups physically separated by environment? Our analysis of genes encoding antibiotic and metal resistance and efflux pumps shows clear differences between subgroups, suggesting the accessory genes of a subgroup may collectively reflect the environment of the bacterial plasmid hosts (Popowska and Krawczyk-Balska 2013; Król et al. 2012; Li et al. 2016). However, it is unclear if this represents a cause or consequence of subgroup diversification. Others have hypothesized that subgroup diversification is driven by plasmid-host specialization (Yano et al. 2013). Unfortunately, only a limited number of examples of plasmid adaption to bacterial hosts currently exist to test this (Bouma and Lenski 1988; Dahlberg and Chao 2003; Stalder et al. 2017; Bottery et al. 2017; Sota et al. 2010; Hughes et al. 2012; Yano et al. 2016; De Gelder et al. 2008). We recently showed that a IncP-1 plasmid gene that imposed a cost to *E. coli* strains and was rapidly lost in these hosts in the laboratory, was, only found in the IncP-1β subgroup, with one exception (Elg et al. 2025). This suggests at least some of the IncP-1β plasmids, namely those with an intact *upf31* (an accessory gene), have not spent considerable time previously in *E. coli* or closely related taxa, unless they accumulated mutations that inactivated the gene. Such differences in host range may well lead to plasmid backbone diversification. Since *upf31* is only found within the IncP-1β subgroup, this might suggest that the gene factored into a more ancient diversification event relative to the diversification of existing subgroups. However, the presence of an intact *upf31* within the phylogeny of the IncP-1 β was paraphyletic (Elg et al. 2025), indicating it may not be actively contributing to diversification within or between established subgroups. Since *upf31* is known to have a very harmful effect on laboratory *E. coli* strains, and many captured plasmids are mated into *E. coli*, it is possible more plasmids contained full-length *upf31* existed but lost the gene in *E. coli* laboratory passage before sequencing. Further discovery of plasmid adaptation to bacterial hosts, and examination of how they correspond to known subgroup lineages, is needed to assess the role of plasmid-bacteria adaptation in IncP-1 plasmid diversification.

Alternatively, it is possible that IncP-1 plasmids from different subgroups regularly co-exist in bacterial communities but are not jointly present within cells long enough for homologous recombination to frequently occur. IncP-1 plasmids belong to the same incompatibility group, which indicates their general inability to co-exist within a cell lineage for multiple generations due to shared replication or partitioning machinery (Novick 1987). Furthermore, all IncP-1 plasmids in our study contain at least two entry exclusion genes, *trbJ* and *trbL*, that lower the frequency of conjugation of another plasmid into the same cell (Garcillán-Barcia and de la Cruz 2008; Cabezón et al. 2015). Thus, the incompatibility between plasmids combined with entry exclusion provides a rational explanation for the low frequency of detectable recombination observed between IncP-1 plasmids backbones. While one study incidentally found some IncP-1 plasmids could co-exist for up to fifty generations (Chikami et al. 1985), we are otherwise unaware of any systematic examination of the time various IncP-1 plasmids can co-exist intra-cellularly. Such a study is warranted, especially if it explored *(1)* the degree of plasmid backbone similarity, and *(2)* the period of intra-cellular co-existence required for recombination to influence plasmid evolution.

Despite a primarily vertical history with strong recombination isolation, we cannot discount that rare recombination events in plasmid backbone genes could meaningfully impact IncP-1 plasmid evolution. Based on data from IncF and IncW plasmids, it has been hypothesized (Fer Andez-Opez et al. 2006; Boyd et al. 1996) that rare recombination events set conjugative plasmids on different evolutionary trajectories leading to diversification. Thus, the rare recombination we see in IncP-1 plasmids might reflect a more generalizable mechanism of plasmid diversification. As shown by Luria and Delbrück in their classic study (Luria and Delbrück 1943), a rare and early evolutionary contingency can have outsized influence on later population genetics (Blount et al. 2018). It is left to be discovered if the recombined *korB* and *kleE* regions found in 22 β-1 plasmids, which appear to be vertically inherited after an initial recombination event, may represent an early subgroup diversification event. More broadly, our predicted regions of recombination and the lineage tree of backbone genes should aid future research into the factors that lead to subgroup diversification of this prevalent and broad-host-range group of plasmids.

## Materials and Methods

### Curation of IncP-1 Dataset and Concatenation of Backbone Genes

The PLSDB (Galata et al. 2019) plasmid database was used along with a literature review to identify IncP-1 plasmid sequences. Our initial set of 206 plasmids was reduced to 187 when partial sequences and those with transposon insertions in backbone genes were removed (Supplemental Data S5).

The number of backbone genes vary by subgroup with an average of 33 genes (Sen et al. 2013). Among our 187 functional plasmid backbone genomes, we identified a set of 20 reliable, conserved backbone genes based on presence and annotation quality that were included in our phylogenetic analyses (Supplemental Data S6). These 20 candidate loci include representatives from the four regions of the plasmids backbone that collectively encode the core functions of IncP-1 plasmids; replication, stable inheritance, control, and the conjugative transfer regions *tra*, and *trb* (Thorsted et al. 1996; Macartney et al. 1997; Figure 1). The replication initiation gene *trfA1* was utilized, as the *ssb* gene was interestingly missing from some plasmids in our dataset. From the central control region, which includes genes for the stable inheritance and regulation of replication and transfer function, we utilized *korB, korA*, and *kleE*. For the *tra* region that encodes genes for processing plasmid transfer DNA we included *traE, F, G, I, J, L*. In the *trb* region that encodes the mating pore and conjugative pilus for plasmid transmission we used *trbA, B, C, D, F, G, H, I, J, L*. These individual gene sequences were aligned using mafft v7.429 (Katoh and Standley 2013). The alignment file for concatenated backbone genes of all 187 plasmids is available in Supplemental Data S7.

### Inference of the IncP-1 phylogeny

We reconstructed individual gene trees for the 187 IncP-1 plasmids for all candidate loci. To distinguish between the random sorting of ancestral alleles (incomplete lineage sorting; ILS) and homologous recombination, we utilized summary-based methods of phylogenetic reconstruction under the multispecies coalescent model (MSC; Pamilo and Nei 1988; Rannala and Yang 2003; Knowles and Carstens 2007). First, we generated maximum-likelihood phylogenies from each of the 20 aligned backbone genes using IQ-TREE v2.2.5 (Minh et al. 2020). Models of nucleotide evolution were selected under Bayesian information criteria (BIC) using ModelFinder (Kalyaanamoorthy et al. 2017) as implemented within IQ-TREE, and maximum-likelihood phylogenies were estimated for each gene independently using 1000 ultra-fast bootstrap replicates (Hoang et al. 2018). From our 20 independent gene tree topologies, we estimated a single lineage tree using ASTRAL-III v5.7.367 (Zhang et al. 2018). The final ASTRAL-III lineage tree was rooted on the IncP-1γ clade as outgroup as done in Sen et al (2013). Our final summary lineage tree was visualized and annotated using the Interactive Tree of Life (Letunic and Bork 2024) and are publicly available to view at https://itol.embl.de/shared/1SsM2kpxu4zAB.

### Detection of Backbone Gene Recombination

We identified regions of the plasmid backbone that are involved in recombination at a nucleotide level using fastGEAR (Mostowy et al. 2017). This program uses a lineage-informed hidden Markov model (HMM) to detect microbial recombination by using polymorphic sites between an individual sequence and other sequences in same lineage to determines whether it is more likely that it originated from an external lineage. The HMM will then determine the alternative lineage from which it was most likely received. We first used fastGEAR’s built-in Bayesian analysis of population structure (BAPS; Corander et al. 2003) to perform lineage clustering, which produced clusters consistent with observed clades in our IncP-1 lineage tree. We then took our IncP-1 clades as illustrated in Figure 2 as inputs for fastGEAR using the ‘current’ setting, resulting in the output for Figure 3 and Supplemental Data S8.

### Comparison of accessory gene patterns between subgroups

Different patterns of resistance gene acquisition between plasmids can contribute to environment specificity potentially leading to the isolation and diversification of plasmid subgroups hosts (Popowska and Krawczyk-Balska 2013; Król et al. 2012; Li et al. 2016). To examine this in our set of 187 IncP-1 plasmid genomes we searched our genomes for genes categorized as *(i)* drugs resistance, *(ii)* metal resistance, and *(iii)* ‘multi-compound’ resistance as classified within the MEGAres database (Bonin et al. 2023). The latter refers to genes like those encoding efflux pumps, which confer at least some resistance to multiple factors (drugs, metals, biocides, *etc*.). Genes within our plasmid genomes that had greater than 90% coverage to gene accessions within the database were identified and returned. The gene accession with the highest coverage was chosen in cases where a plasmid gene had greater than 90% coverage to multiple resistant gene accessions. For the full list of unique gene accessions returned by our search, see Supplemental Data S4. To show differential patterns of resistance gene categories between subgroups, we took the average number of accessory genes for each category found in each subgroup and scaled it to the average number of genes for all plasmids.

## Supporting information

Supplemental Data Description

## Acknowledgements

We are thankful to Olivia Kosterlitz (University of Washington, University of Idaho) and Luke Harmon (University of Idaho) for making important improvements to this manuscript. Thank you to Thibault Stalder (INSERM) for assisting with resistance gene identification. E.M.T. and

C.A.E. received support for this work from National Institute of Allergy and Infectious Diseases Extramural Activities grant no. R01AI084918 from the National Institutes of Health. C.A.E. was supported by the NSF Graduate Research Fellowship (GRFP) grant no. DGE-2019265372.

C.A.E and D.J.S. were both supported by the Bioinformatics and Computational Biology (BCB) Fellowship and the Paul Joyce Memorial BCB Fellowship Endowment at the University of Idaho.

## References

Bahl, M. I., Hansen, L. H., Goesmann, A., & Sørensen, S. J. (2007). The multiple antibiotic resistance IncP-1 plasmid pKJK5 isolated from a soil environment is phylogenetically divergent from members of the previously established α, β and δ sub-groups. Plasmid, 58(1), 31–43.

Bahl, M. I., Burmølle, M., Meisner, A., Hansen, L. H., & Sørensen, S. J. (2009). All IncP-1 plasmid subgroups, including the novel ε subgroup, are prevalent in the influent of a Danish wastewater treatment plant. Plasmid, 62(2), 134–139.

Blount, Z. D., Lenski, R. E., & Losos, J. B. (2018). Contingency and determinism in evolution: Replaying life’s tape. Science, 362(6415), eaam5979.

Bonin, N., Doster, E., Worley, H., Pinnell, L. J., Bravo, J. E., Ferm, P., Marini, S. Prosperi, M., Noyes, N., Morley, P. S., & Boucher, C. (2023). MEGARes and AMR++, v3. 0: an updated comprehensive database of antimicrobial resistance determinants and an improved software pipeline for classification using high-throughput sequencing. Nucleic acids research, 51(D1), D744–D752.

Bottery, M. J., Wood, A. J., & Brockhurst, M. A. (2017). Adaptive modulation of antibiotic resistance through intragenomic coevolution. Nature ecology & evolution, 1(9), 1364–1369.

Bouma, J. E., & Lenski, R. E. (1988). Evolution of a bacteria/plasmid association. Nature, 335(6188), 351–352.

Boyd, E. F., Hill, C. W., Rich, S. M., & Hartl, D. L. (1996). Mosaic structure of plasmids from natural populations of Escherichia coli. Genetics, 143(3), 1091–1100.

Brown, C. J., Sen, D., Yano, H., Bauer, M. L., Rogers, L. M., Van der Auwera, G. A., & Top, E. M. (2013). Diverse broad-host-range plasmids from freshwater carry few accessory genes. Applied and environmental microbiology, 79(24), 7684–7695.

Bruen, T. C., Philippe, H., & Bryant, D. (2006). A simple and robust statistical test for detecting the presence of recombination. Genetics, 172(4), 2665–2681.

Cabezón, E., Ripoll-Rozada, J., Peña, A., De La Cruz, F., & Arechaga, I. (2015). Towards an integrated model of bacterial conjugation. FEMS microbiology reviews, 39(1), 81–95.

Carstens, B. C., & Knowles, L. L. (2007). Estimating species phylogeny from gene-tree probabilities despite incomplete lineage sorting: an example from Melanoplus grasshoppers. Systematic biology, 56(3), 400–411.

Castañeda-Barba, S., Top, E. M., & Stalder, T. (2024). Plasmids, a molecular cornerstone of antimicrobial resistance in the One Health era. Nature Reviews Microbiology, 22(1), 18–32.

Chikami, G. K., Guiney, D. G., Schmidhauser, T. J., & Helinski, D. R. (1985). Comparison of 10 IncP plasmids: homology in the regions involved in plasmid replication. Journal of bacteriology, 162(2), 656–660.

Corander, J., Waldmann, P., & Sillanpää, M. J. (2003). Bayesian analysis of genetic differentiation between populations. Genetics, 163(1), 367–374.

Dahlberg, C., & Chao, L. (2003). Amelioration of the cost of conjugative plasmid carriage in Escherichia coli K12. Genetics, 165(4), 1641–1649.

Datta, N., & Hedges, R. W. (1972). Host ranges of R factors. Microbiology, 70(3), 453–460.

De Gelder, L., Ponciano, J. M., Joyce, P., & Top, E. M. (2007). Stability of a promiscuous plasmid in different hosts: no guarantee for a long-term relationship. Microbiology, 153(2), 452–463.

De Gelder, L., Williams, J. J., Ponciano, J. M., Sota, M., & Top, E. M. (2008). Adaptive plasmid evolution results in host-range expansion of a broad-host-range plasmid. Genetics, 178(4), 2179–2190.

Elg, C. A., Mack, E., Rolfsmeier, M., McLean, T. C., Sneddon, D., Kosterlitz, O., Soderling, E., Narum, S., Rowley, P. A., Sullivan, J., Thomas, C. M., & Top, E. M. (2025) Evolution of a Plasmid Regulatory Circuit Ameliorates Plasmid Fitness Cost. Molecular biology and evolution, 42(4) msaf062.

Fernández-López, R., Garcillán-Barcia, M. P., Revilla, C., Lázaro, M., Vielva, L., & De La Cruz, F. (2006). Dynamics of the IncW genetic backbone imply general trends in conjugative plasmid evolution. FEMS microbiology reviews, 30(6), 942–966.

Frost, L. S., Leplae, R., Summers, A. O., & Toussaint, A. (2005). Mobile genetic elements: the agents of open source evolution. Nature Reviews Microbiology, 3(9), 722–732.

Galata, V., Fehlmann, T., Backes, C., & Keller, A. (2019). PLSDB: a resource of complete bacterial plasmids. Nucleic acids research, 47(D1), D195–D202.

Garcillán-Barcia, M. P., & de la Cruz, F. (2008). Why is entry exclusion an essential feature of conjugative plasmids? Plasmid, 60(1), 1–18.

Goldenfeld, N., & Woese, C. (2007). Biology’s next revolution. Nature, 445(7130), 369.

Haines, A. S., Akhtar, P., Stephens, E. R., Jones, K., Thomas, C. M., Perkins, C. D., Williams, J. R., Day, M. J., & Fry, J. C. (2006). Plasmids from freshwater environments capable of IncQ retrotransfer are diverse and include pQKH54, a new IncP-1 subgroup archetype. Microbiology, 152(9), 2689–2701.

Hall, J. P., Brockhurst, M. A., & Harrison, E. (2017). Sampling the mobile gene pool: innovation via horizontal gene transfer in bacteria. Philosophical Transactions of the Royal Society B: Biological Sciences, 372(1735), 20160424.

Hayakawa, M., Tokuda, M., Kaneko, K., Nakamichi, K., Yamamoto, Y., Kamijo, T., Umeki, H., Chiba, R., Yamada, R., Mori, M., Yanagiya, K., Moriuchi, R., Yuki, M., Dohra, H., Futamata, H., Ohkuma, M., Kimbara, K., & Shintani, M. (2022). Hitherto-unnoticed self-transmissible plasmids widely distributed among different environments in Japan. Applied and Environmental Microbiology, 88(18), e01114–22.

Heinemann, J. A., & Sprague Jr, G. F. (1989). Bacterial conjugative plasmids mobilize DNA transfer between bacteria and yeast. Nature, 340(6230), 205–209.

Hill, K. E., Weightman, A. J., & Fry, J. C. (1992). Isolation and screening of plasmids from the epilithon which mobilize recombinant plasmid pD10. Applied and Environmental Microbiology, 58(4), 1292–1300.

Hoang, D. T., Chernomor, O., Von Haeseler, A., Minh, B. Q., & Vinh, L. S. (2018). UFBoot2: improving the ultrafast bootstrap approximation. Molecular biology and evolution, 35(2), 518–522.

Hughes, J. M., Lohman, B. K., Deckert, G. E., Nichols, E. P., Settles, M., Abdo, Z., & Top, E. M. (2012). The role of clonal interference in the evolutionary dynamics of plasmid-host adaptation. MBio, 3(4), 10–1128.

Kalyaanamoorthy, S., Minh, B. Q., Wong, T. K., Von Haeseler, A., & Jermiin, L. S. (2017). ModelFinder: fast model selection for accurate phylogenetic estimates. Nature methods, 14(6), 587–589.

Katoh, K., & Standley, D. M. (2013). MAFFT multiple sequence alignment software version 7: improvements in performance and usability. Molecular biology and evolution, 30(4), 772–780.

Klümper, U., Riber, L., Dechesne, A., Sannazzarro, A., Hansen, L. H., Sørensen, S. J., & Smets, B. F. (2015). Broad host range plasmids can invade an unexpectedly diverse fraction of a soil bacterial community. The ISME journal, 9(4), 934–945.

Kottara, A., Hall, J. P., Harrison, E., & Brockhurst, M. A. (2016). Multi-host environments select for host-generalist conjugative plasmids. BMC Evolutionary Biology, 16, 1–4.

Król, J. E., Penrod, J. T., McCaslin, H., Rogers, L. M., Yano, H., Stancik, A. D., Dejonghe, W., Brown, C. J., Parales, R. E., Wuertz, S., & Top, E. M. (2012). Role of IncP-1β plasmids pWDL7:: rfp and pNB8c in chloroaniline catabolism as determined by genomic and functional analyses. Applied and environmental microbiology, 78(3), 828–838.

Letunic, I., & Bork, P. (2021). Interactive Tree Of Life (iTOL) v5: an online tool for phylogenetic tree display and annotation. Nucleic acids research, 49(W1), W293–W296.

Li, X., Wang, Y., Brown, C. J., Yao, F., Jiang, Y., Top, E. M., & Li, H. (2016). Diversification of broad host range plasmids correlates with the presence of antibiotic resistance genes. FEMS microbiology ecology, 92(1), fiv151.

Luria, S. E., & Delbrück, M. (1943). Mutations of bacteria from virus sensitivity to virus resistance. Genetics, 28(6), 491.

Macartney, D. P., Williams, D. R., Stafford, T., & Thomas, C. M. (1997). Divergence and conservation of the partitioning and global regulation functions in the central control region of the IncP plasmids RK2 and R751. Microbiology, 143(7), 2167–2177.

Maddison, W. P. (1997). Gene trees in species trees. Systematic biology, 46(3), 523–536.

Minh, B. Q., Schmidt, H. A., Chernomor, O., Schrempf, D., Woodhams, M. D., Von Haeseler, A., & Lanfear, R. (2020). IQ-TREE 2: new models and efficient methods for phylogenetic inference in the genomic era. Molecular biology and evolution, 37(5), 1530–1534.

Mostowy, R., Croucher, N. J., Andam, C. P., Corander, J., Hanage, W. P., & Marttinen, P. (2017). Efficient inference of recent and ancestral recombination within bacterial populations. Molecular biology and evolution, 34(5), 1167–1182.

Musovic, S., Oregaard, G., Kroer, N., & Sørensen, S. J. (2006). Cultivation-independent examination of horizontal transfer and host range of an IncP-1 plasmid among gram-positive and gram-negative bacteria indigenous to the barley rhizosphere. Applied and environmental microbiology, 72(10), 6687–6692.

Norberg, P., Bergström, M., & Hermansson, M. (2014). Complete nucleotide sequence and analysis of two conjugative broad host range plasmids from a marine microbial biofilm. PLoS One, 9(3), e92321.

Norberg, P., Bergström, M., Jethava, V., Dubhashi, D., & Hermansson, M. (2011). The IncP-1 plasmid backbone adapts to different host bacterial species and evolves through homologous recombination. Nature communications, 2(1), 268.

Norman, A., Hansen, L. H., & Sørensen, S. J. (2009). Conjugative plasmids: vessels of the communal gene pool. Philosophical Transactions of the Royal Society B: Biological Sciences, 364(1527), 2275–2289.

Novick, R. P. (1987). Plasmid incompatibility. Microbiological reviews, 51(4), 381–395.

Olsen, R. H., & Shipley, P. (1973). Host range and properties of the Pseudomonas aeruginosa R factor R1822. Journal of Bacteriology, 113(2), 772–780.

Pamilo, P., & Nei, M. (1988). Relationships between gene trees and species trees. Molecular biology and evolution, 5(5), 568–583.

Pansegrau, W., Lanka, E., Barth, P. T., Figurski, D. H., Guiney, D. G., Haas, D., Helinski, D. R., Schwab, H., Stanisich, V. A., & Thomas, C. M. (1994). Complete nucleotide sequence of Birmingham IncPα plasmids: compilation and comparative analysis. Journal of molecular biology, 239(5), 623–663.

Partridge, S. R., Kwong, S. M., Firth, N., & Jensen, S. O. (2018). Mobile genetic elements associated with antimicrobial resistance. Clinical microbiology reviews, 31(4), 10–1128.

Pilla, G., & Tang, C. M. (2018). Going around in circles: virulence plasmids in enteric pathogens. Nature Reviews Microbiology, 16(8), 484–495.

Popowska, M., & Krawczyk-Balska, A. (2013). Broad-host-range IncP-1 plasmids and their resistance potential. Frontiers in microbiology, 4, 44.

Rannala, B., & Yang, Z. (2003). Bayes estimation of species divergence times and ancestral population sizes using DNA sequences from multiple loci. Genetics, 164(4), 1645–1656.

Schlüter, A., Heuer, H., Szczepanowski, R., Forney, L. J., Thomas, C. M., Puhler, A., & Top, E. M. (2003). The 64 508 bp IncP-1 β antibiotic multiresistance plasmid pB10 isolated from a waste-water treatment plant provides evidence for recombination between members of different branches of the IncP-1 β group. Microbiology, 149(11), 3139–3153.

Schlüter, A., Szczepanowski, R., Pühler, A., & Top, E. M. (2007). Genomics of IncP-1 antibiotic resistance plasmids isolated from wastewater treatment plants provides evidence for a widely accessible drug resistance gene pool. FEMS microbiology reviews, 31(4), 449–477.

Schmidhauser, T. J., & Helinski, D. R. (1985). Regions of broad-host-range plasmid RK2 involved in replication and stable maintenance in nine species of gram-negative bacteria. Journal of bacteriology, 164(1), 446–455.

Sen, D., Brown, C. J., Top, E. M., & Sullivan, J. (2013). Inferring the evolutionary history of IncP-1 plasmids despite incongruence among backbone gene trees. Molecular biology and evolution, 30(1), 154–166.

Sen, D., Van der Auwera, G. A., Rogers, L. M., Thomas, C. M., Brown, C. J., & Top, E. M. (2011). Broad-host-range plasmids from agricultural soils have IncP-1 backbones with diverse accessory genes. Applied and environmental microbiology, 77(22), 7975–7983.

Sen, D., Yano, H., Suzuki, H., Król, J. E., Rogers, L., Brown, C. J., & Top, E. M. (2010). Comparative genomics of pAKD4, the prototype IncP-1δ plasmid with a complete backbone. Plasmid, 63(2), 98–107.

Shintani, M., Takahashi, Y., Yamane, H., & Nojiri, H. (2010). The behavior and significance of degradative plasmids belonging to Inc groups in Pseudomonas within natural environments and microcosms. Microbes and environments, 25(4), 253–265.

Sota, M., Tsuda, M., Yano, H., Suzuki, H., Forney, L. J., & Top, E. M. (2007). Region-specific insertion of transposons in combination with selection for high plasmid transferability and stability accounts for the structural similarity of IncP-1 plasmids. Journal of bacteriology, 189(8), 3091–3098.

Sota, M., Yano, H., M Hughes, J., Daughdrill, G. W., Abdo, Z., Forney, L. J., & Top, E. M. (2010). Shifts in the host range of a promiscuous plasmid through parallel evolution of its replication initiation protein. The ISME journal, 4(12), 1568–1580.

Soucy, S. M., Huang, J., & Gogarten, J. P. (2015). Horizontal gene transfer: building the web of life. Nature Reviews Genetics, 16(8), 472–482.

Stalder, T., Rogers, L. M., Renfrow, C., Yano, H., Smith, Z., & Top, E. M. (2017). Emerging patterns of plasmid-host coevolution that stabilize antibiotic resistance. Scientific Reports, 7(1), 4853.

Swenson, N. G. (2009). Phylogenetic resolution and quantifying the phylogenetic diversity and dispersion of communities. PloS one, 4(2), e4390.

Thomas, C. M., & Smith, C. A. (1987). Incompatibility group P plasmids: genetics, evolution, and use in genetic manipulation. Annual Reviews in Microbiology, 41(1), 77–101.

Thorsted, P. B., Shah, D. S., Macartney, D., Kostelidou, K., & Thomas, C. M. (1996). Conservation of the genetic switch between replication and transfer genes of IncP plasmids but divergence of the replication functions which are major host-range determinants. Plasmid, 36(2), 95–111.

Vedler, E., Vahter, M., & Heinaru, A. (2004). The completely sequenced plasmid pEST4011 contains a novel IncP1 backbone and a catabolic transposon harboring tfd genes for 2, 4-dichlorophenoxyacetic acid degradation. Journal of bacteriology, 186(21), 7161–7174.

Yakimov, M. M., Crisafi, F., Messina, E., Smedile, F., Lopatina, A., Denaro, R., Pieper, D. H., Galyshin, P. N., & Giuliano, L. (2016). Analysis of defence systems and a conjugative IncP-1 plasmid in the marine polyaromatic hydrocarbons-degrading bacterium Cycloclasticus sp. 78-ME. Environmental Microbiology Reports, 8(4), 508–519.

Yano, H., Deckert, G. E., Rogers, L. M., & Top, E. M. (2012). Roles of long and short replication initiation proteins in the fate of IncP-1 plasmids. Journal of bacteriology, 194(6), 1533–1543.

Yano, H., Wegrzyn, K., Loftie-Eaton, W., Johnson, J., Deckert, G. E., Rogers, L. M., Konieczny, I. K., & Top, E. M. (2016). Evolved plasmid-host interactions reduce plasmid interference cost. Molecular microbiology, 101(5), 743–756.

Yano, H., Rogers, L. M., Knox, M. G., Heuer, H., Smalla, K., Brown, C. J., & Top, E. M. (2013). Host range diversification within the IncP-1 plasmid group. Microbiology, 159(Pt_11), 2303–2315.

Zhang, C., Rabiee, M., Sayyari, E., & Mirarab, S. (2018). ASTRAL-III: polynomial time species tree reconstruction from partially resolved gene trees. BMC bioinformatics, 19, 15–30.

