## Supplemental Data Description for "Not So Mosaic After All? The Core Genomes of IncP-1 Plasmids Evolve Predominantly Through Vertical Transmission"

**Supplemental Data S1:** All individual backbone gene phylogenies

**Supplemental Data S2:** Individual backbone gene phylogeny topology classifications based on Sen et al. 2013

**Supplemental Data S3:** High resolution version of main text Figure 3

**Supplemental Data S4:** Complete list of drug, metal, and multi-compound resistance gene accessions returned from MEGAres database

**Supplemental Data S5:** Full list of plasmids used in phylogenetic analyses

**Supplemental Data S6:** Backbone gene fasta files

**Supplemental Data S7:** Concatenated plasmid backbone alignment

**Supplemental Data S8:** Full output from fastGEAR
