## Supplementary figures and images for "Not So Mosaic After All? The Core Genomes of IncP-1 Plasmids Evolve Predominantly Through Vertical Transmission"

### Supplemental Data S3

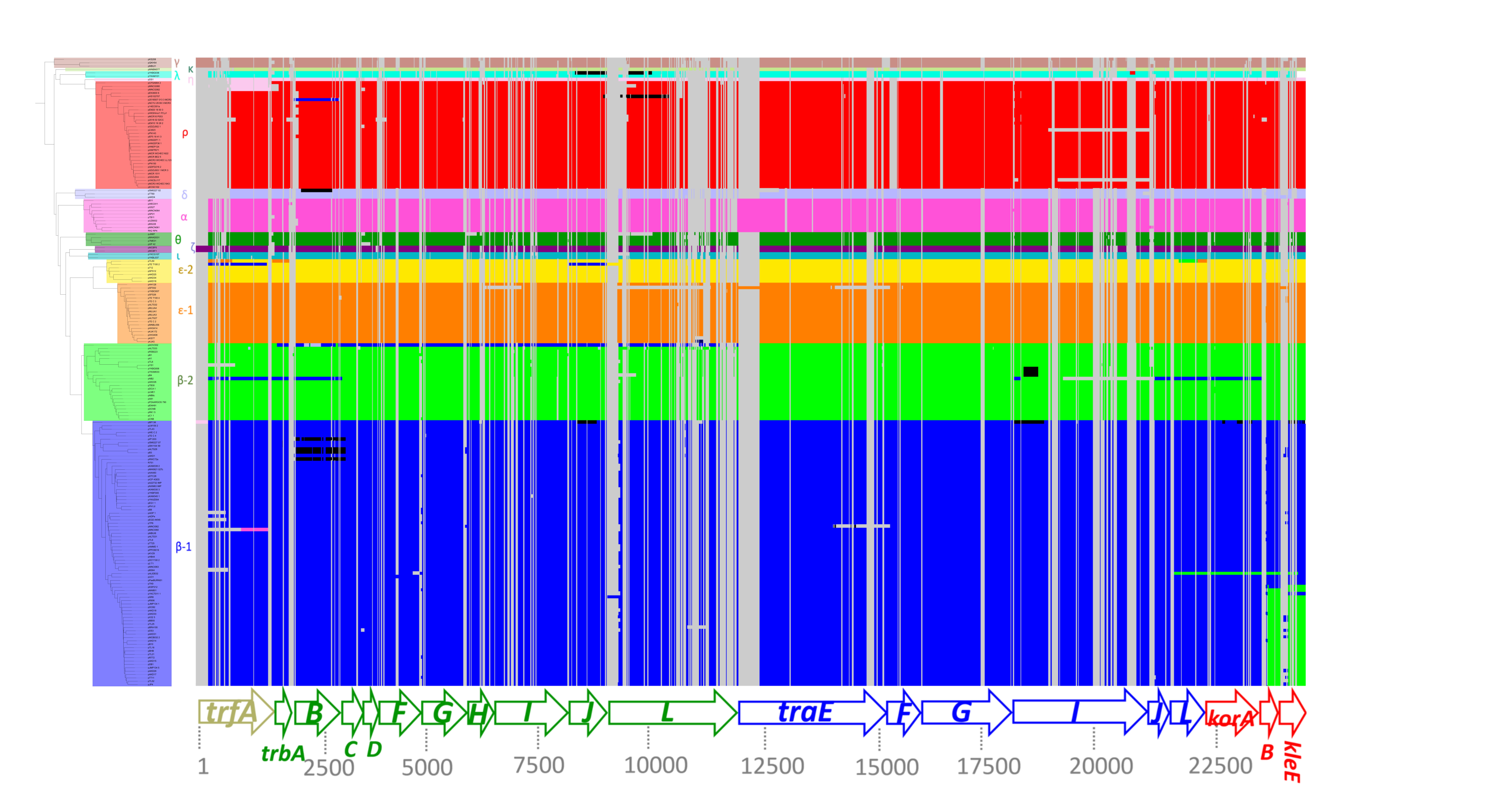
